# Rational Design of RNA Structures that Predictably Tune Eukaryotic Gene Expression

**DOI:** 10.1101/137877

**Authors:** Tim Weenink, Robert M. McKiernan, Tom Ellis

## Abstract

Predictable tuning of gene expression is essential for engineering genetic circuits and for optimising enzyme levels in metabolic engineering projects. In bacteria, gene expression can be tuned at the stage of transcription, by exchanging the promoter, or at stage of translation by altering the ribosome binding site sequence. In eukaryotes, however, only promoter exchange is regularly used, as the tools to modulate translation are lacking. Working in *S. cerevisiae* yeast, we here describe how hairpin RNA structures inserted into the 5’ untranslated region (5’UTR) of mRNAs can be used to tune expression levels by altering the efficiency of translation initiation. We demonstrate a direct link between the calculated free energy of folding in the 5’UTR and protein abundance, and show that this enables rational design of hairpin libraries that give predicted expression outputs. Our approach is modular, working with different promoters and protein coding sequences, and it outperforms promoter mutation as a way to predictably generate a library where a protein is induced to express at a range of different levels. With this tool, computational RNA sequence design can be used to predictably fine-tune protein production, providing a new way to modulate gene expression in eukaryotes.

## Introduction

The rate of production of a given protein in a cell is determined by its gene expression, which in its most simple form is a two-step process, with the gene first transcribed into an mRNA as directed by the promoter, and this mRNA then translated by the ribosome to make the protein. Altering the expression levels of genes is a crucial tool for modern bioscience research, for synthetic biology and for many biotechnology applications [1, 2, 3], and in all organisms this is most-readily achieved by intervening in the first step of gene expression, by changing or regulating the promoter in order to change the rate of transcription of the mRNA [4, 5]. Natural systems, however, also regularly modify protein production by altering the second stage of gene expression where the mRNA is translated [6]. In model bacterial systems, changing the gene sequence in order to modify the rate of translation is a well-established method for predictably tuning the rate of protein production from a gene. However, this is a rarely-used strategy for engineering changes in gene expression in eukaryotic systems.

In bacteria the rate of translation of an mRNA and thus the expression level of a gene can be tuned by changing the sequence of bases immediately upstream of the AUG start codon that are known to recruit the ribosome to initiate translation through RNA:RNA base-pairing [7]. This region, known as the Ribosome Binding Site (RBS) consists of a core sequence that directly base-pairs with the 16S RNA of the ribosome, and surrounding sequences that modify the efficiency of this interaction by forming local secondary structures via mRNA folding. As the efficiency of ribosome recruitement at the RBS defines the translation initiation rate of an mRNA, extensive research has been undertaken to determine how changes to the RBS sequence can be used to tune gene expression [8]. This has led to several sequence-to-output predictive tools that use thermodynamic models of nucleic acid pairing to predict the binding efficiency of ribosomes to any given bacterial mRNA [7]. These are enabled by the multiple software packages that predict nucleic acid secondary structures and determine their Minimum Free Energy (MFE) of folding by summing the thermodynamic contributions of all base-pairing interactions [9, 10, 11, 12, 13, 14].

The most advanced RBS prediction tool, the RBS Calculator, uses secondary-structure calculations to predict the strengths of the various RNA:RNA interactions that occur during ribosome binding and converts these into a predicted translation rate [8]. It can also forward-design new 5’UTR sequences in order to produce desired gene expression levels and can design libraries of 5’UTR sequences in which the careful placement of a few degenerate bases leads to a diverse, yet bounded, variation in the resulting expression levels from the library of DNA expression constructs that encodes these [8, 15]. This enables researchers working with bacteria to be able to fine-tune the expression of genes of interest and design graded expression libraries in a predictable manner, greatly accelerating progress in synthetic biology and biotechnology applications such as metabolic engineering.

In eukaryotes, translation initiation follows a different mechanism to bacteria, where only part of the ribosome, the 40S subunit, initially binds the mRNA [16]. After binding the 5’ cap, it scans along the 5’UTR of the mRNA until reaching the first AUG start codon, which is usually preceded by an A or G/C rich upstream motif known as the Kozak sequence [17]. No direct RNA:RNA base pairing is seen between the ribosome and the mRNA, and as such changing the bases within this region is a rarely-used mechanism for altering gene expression.

However, as with bacteria, it is well-established that the folding of secondary structures in the mRNA 5’UTR can affect the rate of translation and typically when this occurs it inhibits protein expression [18, 19, 20]. Three genome-wide studies in *S. cerevisiae* have shown that a negative correlation exists between the efficiency with which an mRNA is translated and the secondary structure around its start codon [21, 22, 23], and in a recent study where the 10 bases upstream of the start codon on an mRNA were randomised, a significant association between thermodynamically stable secondary structures and reduced protein levels was found [24]. Secondary structures present in 5’UTRs in higher eukaryotes, such as mammalian cells [25, 26] and plants [27], have also been shown to lead to a similar reduction in gene expression.

In several past experiments in yeast, secondary structures have been added to mRNAs to regulate and tune their expression. Lamping *et al.* recently showed that GC-rich sequences encoding hairpins can be used as modules to down-regulate expression [20], while others have combined hairpins with other forms of RNA-based translational regulation or with RNA-binding proteins that bind these motifs [28, 29, 30, 31]. No approach, however, exists where gene expression can be predictably tuned at the translation step in eukaryotes as it can in bacteria. Achieving tools equivalent to the RBS Calculator in eukaryotes like yeast would be highly-desirable, especially for metabolic engineering projects where enzyme expression levels need to be optimised, or in synthetic biology where precise and efficient gene expression is typically desired [2, 3].

Towards this objective, we describe here a design-led approach to predictably tune the expression of genes in *S. cerevisiae* by repressing translation of mRNAs through the introduction of synthetic hairpin secondary structures within the 5’UTR. Following the strategy established by the RBS Library Designer tool [15], we use MFE-based prediction of RNA secondary structures to design sequences with degenerate bases that can be inserted into the 5’UTR to yield a defined range of expression levels within a population of yeast. We derive a mathematical model that links the predicted MFE of folding for each hairpin sequence with resulting protein expression levels measured *in vivo* and show that sequences can be designed to fine-tune gene expression libraries as desired. We show that these hairpin-encoding sequences are modular, working as predicted when paired with different promoters or alternative protein-coding sequences, and are able to alter expression from regulated promoters without impairing their regulation. Our approach greatly simplifies the production of graded expression libraries in *S. cerevisiae* compared to existing methods and offers a valuable new tool for predictably tuning gene expression in eukaryotes.

## Materials & Methods

### Strains and media

BY4741 (MATa his3Δ1 leu2Δ0 met15Δ0 ura3Δ0) was used for all yeast transformations, using a high efficiency yeast transformation protocol [32]. Standard practice in yeast genetics was followed [33]. When used, IPTG was supplied at a concentration of 10mM. For selection in *E. coli*, antibiotics were added at the following concentrations: Ampicillin: 100 *μ*g/ml, Kanamycin: 50 *μ*g/ml, Chloramphenicol: 33 *μ*g/ml.

### Library cloning

The Yeast ToolKit (YTK) cloning system was used for library plasmid construction [34]. NEB Turbo chemically competent *E. coli* (Catalog #C2984I) were used for transformation of library constructs, according to the manufacturer’s protocol. Supplementary table 1 lists all multigene level plasmids constructed for the libraries in this study and their constituent cassette-level plasmids. Contrary to normal YTK protocol, one of the cassettes is an *in vitro* product created via PCR, rather than through a cloning step. The corresponding PCR reactions for each cassette part in these libraries are also shown in Supplementary table 1 and primers are defined in Supplementary table 4. For a detailed description of the cloning method we refer to the supplementary material and Supplementary figure 1.

Cloned cassette-level plasmids used in the multigene assembly reactions and as template for the PCRs are listed in Supplementary table 2, along with part plasmids used in their assemblies. Any used parts that were not defined as a standard part in the YTK kit are listed in Supplementary table 3.

### Flow cytometry analysis

Libraries were tested using flow cytometry performed with the Attune NxT Acoustic Focusing Cytometer (Invitrogen), with accompanying 96-well plate reader. Two lasers are installed in this machine: blue 488 nm and a 561 nm yellow laser. Green fluorescence (blue laser) was detected at a voltage of 450, red fluorescence (yellow laser) at a voltage of 480, while forward and side scatter were detected at voltages of 40 and 340, respectively.

Strains were picked and grown to saturation in 700*μ*l YEP-Dextrose in 2ml 96-deepwell plates (VWR, cat no 732-0585). Cultures were grown overnight in a shaking incubator (Infors HT multitron MTP shaker) at 800rpm at 30°C, with breathe-easy film (sigma, cat no Z380059) covering the plate to prevent evaporation. This plate was then diluted 500 times into a new deepwell plate containing 700*μ*l minimal dropout media with 2% galactose for induction. After overnight incubation of at least 12 hours, the cultures were backdiluted into a Costar 96 round-well flatbottom plate (VWR, cat no 3596). Dilution was 10-100 fold, depending on culture density, using the same media in a total volume of 300*μ*l per well. Cultures were grown for a minimum of 4 hours before initiation of the measurements, to ensure logarithmic growth. For autofluorescence measurements the same incubation protocol was followed, exclusively with dextrose as the carbon source.

10,000 events were collected for each of the 3 biological repeats per sample. Only events with forward and side scatter values greater than 10^3^ were counted. Populations were tightly gated around the median of forward-and side-scatter, in order to limit the effect of cell size on the measurements. Populations were subsequently gated for sufficient mRuby2 expression in strains that contained a constitutively expressed red fluorescent control. Conversely, in constructs with constitutively expressed yEGFP, populations were gated for sufficient green fluorescence. Gating and exporting was done using FlowJo 10.0.7r2. We defined normalised fluorescence as the fold increase over median autofluorescence levels. Accordingly, raw fluorescence values of tested strains were divided by the median fluorescence of unmodified BY4741 cells, grown under identical conditions. Matlab 2016b was used for the visualisation of the resulting histograms.

### Library member sequence determination

Hairpin sequences of individual library members were determined through yeast colony PCR. Single colonies were resuspended in 50 *μ*l 0.02 M NaOH with a sterile toothpick. After incubating this solution at 99°C for 10 minutes, a 2 *μ*l aliquot was used as template in a 50 *μ*l PCR reaction. Primers TW149 and TW188 were used for the reaction and for sequencing. They are defined in Supplementary table 4. Isolates with DNA sequences that did not match the relevant library sequence space (i.e. mutations or sequencing errors) were discarded from the analysis. 5’UTR sequences of all isolated library members are listed in Supplementary table 6.

### Reverse transcription and qPCR

Total RNA was isolated from yeast using the YeaStar RNA Kit (Zymo Research, Cat No R1002), according to the manufacturer’s instructions. Cultures were grown to saturation overnight and backdiluted 1:100 the next morning. Cultures were then grown to an O.D._600_ of approximately 2, to ensure logarithmic growth. 1.5 ml of the culture was used for RNA isolation. This volume was adjusted to ensure that the same amount of biomass was used for every sample.

400 ng of total RNA of each of the samples was used in the reverse transcription (RT) reaction to produce cDNA that could be used for qPCR. The RT reaction was performed in a total volume of 10*μ*l, using the Tetro cDNA synthesis kit (Bioline, Cat No BIO-65043) according to the manufacturer’s instructions. For each reaction, a negative control lacking the reverse transcriptase was included. Specific primers were used for the RT step, which are listed in Supplementary table 4.

cDNA obtained in the RT reaction was diluted 300x and used for qPCR. 4.6*μ*l of diluted cDNA was used in a total reaction volume of 10*μ*l. The Kapa universal qPCR 2x mastermix kit (KAPA biosystems, Cat No kk4601) was used according to the manufacturer’s instructions. The primers (0.2*μ*l per primer per reaction) for each of the screened targets are listed in Supplementary table 4. Measurements were performed with the Eppendorf MasterCycler RealPlex qPCR thermocycler and accompanying software. The following cycling program was used: denaturation for 10 min at 95°C followed by 50 cycles of 15 s at 95°C, 1 min annealing and extension at 60°C.

Three technical replicates were performed for every biological sample. The data were analysed using the 2^-ΔΔ*C*_*T*_^ (also ‘dd-Ct’) method [35]. TPI1 was used as a reference gene [36]. The error was calculated as the standard deviation of the replicates with propagation of the error in the reference gene measurements. For each qRT-PCR experiment, two controls were included to monitor the level of DNA contamination of the cDNA and the used reagents. For every target a triplicate measurement of ddH_2_O was included. Secondly, for every target in every strain we included the -RT control samples produced during the cDNA synthesis.

## Results

### Altering expression strength with 5’UTR RNA structures

We set out to build on recent work in *S. cerevisiae* showing that hairpin structures within the 5’UTR of mRNAs decrease expression by inhibiting translation [20]. To verify this finding and explore the relationship in more depth, we used one-pot cloning methods to introduce a library of hairpin structures into the 5’UTR of an mRNA encoding expression of green fluorescent protein (GFP). This mRNA is expressed from the GAL1 promoter and upon galactose induction yields a strong and easily-detectable green fluorescent signal from yeast cells. If RNA structures placed within the 5’UTR of this mRNA lead to changes in the efficiency of its translation, then the fluorescent signal of the yeast will be altered and the expression level can be quantified by analytical flow cytometry.

Using a strategy of PCR amplification with degenerate oligonucleotide primers, followed by cloning with the modular YTK system, we introduced a hairpin library spaced 15 bases upstream of the start codon of GFP, as summarised in Figure 1 and detailed in Supplementary figure 1. To design the hairpin-encoding sequences, we used RNAfold from the ViennaRNA suite of tools, as it allows the RNA-folding algorithms to run on a local computer and the thermodynamic parameters can be adjusted to work for 30°C [37, 14]. As an initial test, two pairs of degenerate oligonucleotide primers were designed to create two alternative secondary structure libraries. The primer pairs each encoded the introduction of 60 bases of sequence predicted by RNAfold to fold into a strong hairpin motif consisting of a 27 basepair stem and 4 base loop (Figure 1A).

**Figure 1:**
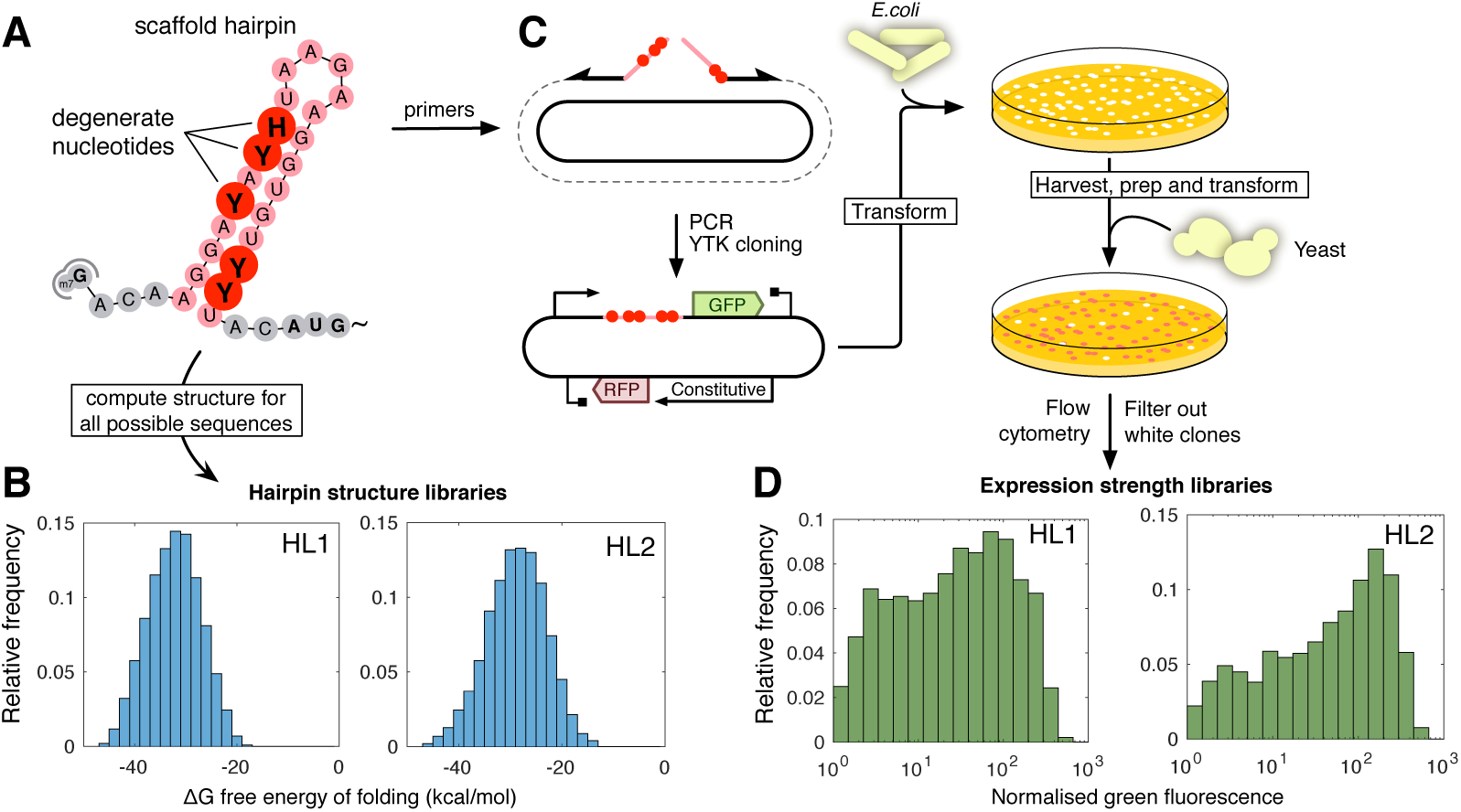
Overview of 5’UTR hairpin library HL1 and HL2 creation. (**A**) Degenerate nucleotides are inserted at various positions into the design of a hairpin scaffold. (**B**) The minimum free energy of all possible sequences is calculated with RNAfold and visualised in a histogram. (**C**) When the design meets the requirements, primers incorporating the required degeneracies are ordered. A library of plasmids for *E. coli* transformation is created using PCR and a subsequent Golden Gate based assembly step implemented in Yeast ToolKit (YTK) format. The library of *E. coli* transformants is harvested and plasmids prepped for yeast transformation. (**D**) Transformant yeast colonies are pooled and analysed using flow cytometry. Clones that do not show constitutive red fluorescence are discarded in quality control for correct assembly. The diversity of the library of hairpins is reflected in the spread of green fluorescence over three orders of magnitude.

Degenerate nucleotides were designed into 10 positions within the primers so that the different constructs produced by the cloning would have variation in the bases within the hairpin stem, and therefore a range of strengths for the resulting RNA secondary structure. Using RNAfold, the distribution of the predicted minimum free energies of folding for each of the two hairpin libraries could be determined by calculating the folding strengths for all possible combinations of introduced degenerate bases (Figure 1B). The two initial libraries, each with 10 degenerate bases were designed to give a normal distribution of predicted secondary structure strengths with average minimum free energy (MFE) of folding of -32.2 kcal/mol and -28.8 kcal/mol (libraries HL1 and HL2, respectively - see Supplementary table 5 for sequences).

Following plasmid construction to introduce the hairpin-encoding sequences with degenerate bases, *E. coli* were transformed and all resulting colonies were pooled for each library. Plasmid libraries were extracted from these two pools and then transformed into *S. cerevisiae* cells. All yeast colonies from each library were pooled and then grown in galactose media to induce gene expression (Figure 1C).

The green fluorescence from each induced pool of yeast colonies was measured at the single-cell level by flow cytometry and used to quantify GFP expression from the library constructs. For both HL1 and HL2 libraries, the normalised green fluorescence was seen to vary across the population over three orders of magnitude (Figure 1D), indicating that the introduced hairpin sequences were indeed altering the expression of GFP in the cells. The shape and peak of the fluorescence histograms in the two cases also differed, with more cells exhibiting low amounts of GFP expression when the library with stronger predicted secondary structure (HL1) was used.

### Matching expression levels to predicted folding energies

We next sought to determine if there was a mathematical relationship between the predicted MFE of the encoded 5’UTR secondary structures and the resulting *in vivo* GFP expression levels in *S. cerevisiae*To do this we selected 31 individual colonies from libraries HL1 and HL2 (Figure 1), determined the sequence of their 5’UTR regions and used flow cytometry to characterise their green fluorescence per cell upon galactose induction. We used RNAfold to predict the MFE for the hairpin structures in each isolate by inputting their 5’UTR sequences and plotted the relationship between the predicted MFE and the normalised green fluorescence measurements (Figure 2A).

**Figure 2:**
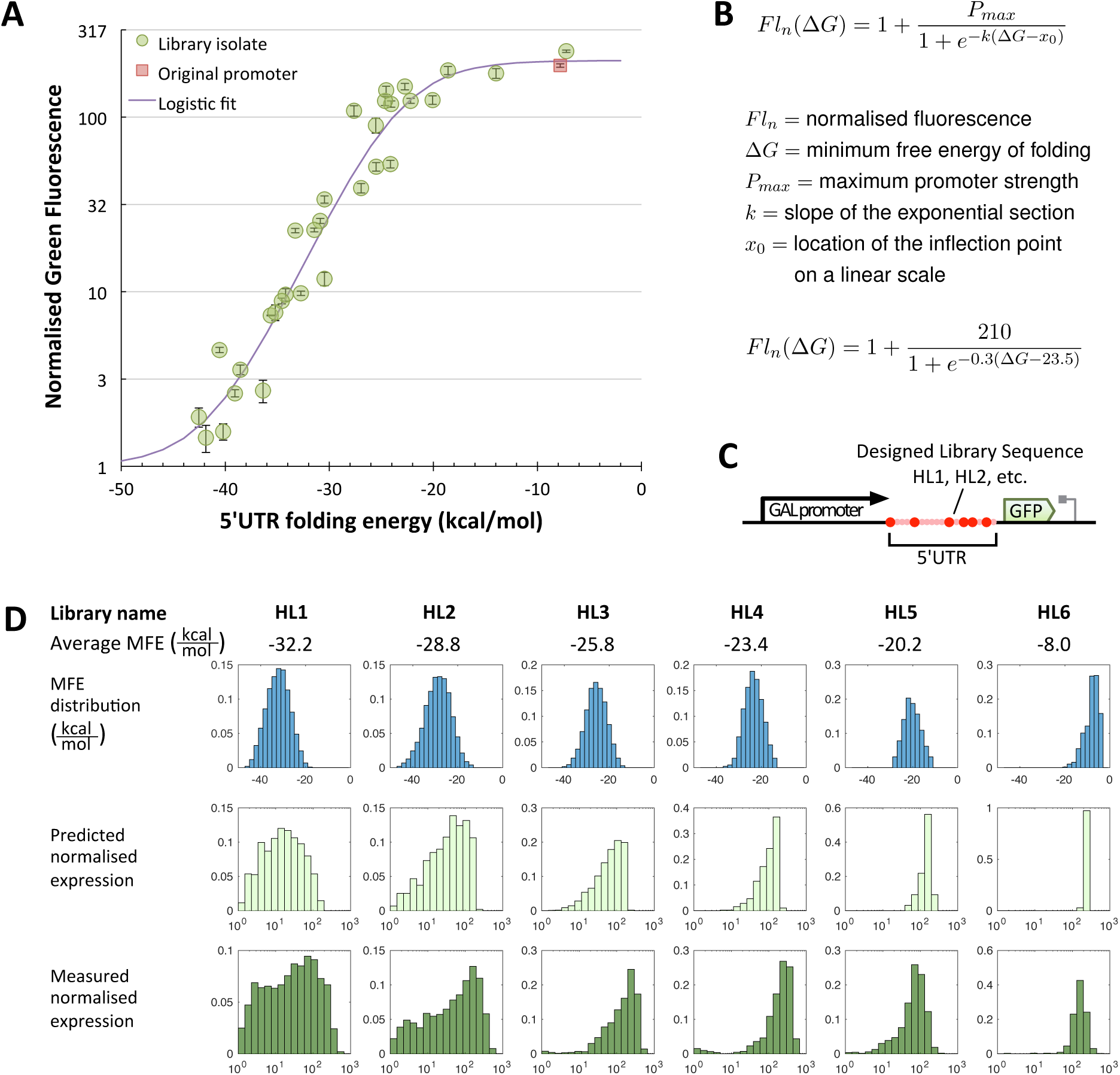
Correlation between 5’UTR hairpin strength and protein expression. Stronger hairpins are shown to cause lower expression in a predictable manner. (**A**) Determination of the transfer function between hairpin folding energy and normalised green fluorescence. Fluorescence was measured for 31 isolated HL1 and HL2 library members and divided by the median autofluorescence of the parental strain to obtain the normalised fluorescence. The isolates were sequenced to obtain the 5’UTR sequences, which were used to calculate the corresponding hairpin folding energy. The diagram shows the folding energy of the 5’UTR of each isolate plotted against the normalised green fluorescence and fitted to a logistic growth curve. Sequences and obtained values are listed in Supplementary table 6. Error bars indicate standard deviation of the median of three measurements of 10,000 events each. (**B**) Equation describing the logistic fit between predicted folding energy and normalised fluorescence. (**C**) Diagram of the transcription unit that constitutes the RNA hairpin library. Pink area constitutes the hairpin backbone with red spheres indicating degenerate nucleotides. (**D**) Correlation between the predicted strength of 5’UTR structure libraries and the measured gene expression distributions of these libraries. All panels show normalised frequency distributions (histograms). A total of 6 libraries (HL1-6) are shown, whose average minimum free energy (MFE) of folding is given in kcal/mol. In the top row of panels, the histogram of the distribution of the MFE in the 5’UTRs of the different libraries is shown. The horizontal axes for these panels ranges on a linear scale from -50 kcal/mol to 0 kcal/mol. The middle row converts these into a histogram of predicted normalised expression levels using the equation established in sub-figure A. The third row shows the experimentally obtained distribution of normalised fluorescent reporter expression levels as measured by flow cytometry. In the lower two rows, the horizontal axis corresponds to normalised green fluorescence (a unit-less quantity) ranging on a logarithmic scale from 1 on the left to 1000 on the right.

This plot revealed a clear relationship between the predicted UTR folding energy and the GFP ex-pression for each isolated colony. A steep decline in the expression level is seen as the MFE approaches more negative values, in line with the hypothesis that increasingly strong secondary structures within the 5’UTR inhibit gene expression. However, this decline does not start until the MFE reaches approximately -22 kcal/mol, which indicates that the structures weaker than -22 kcal/mol are not strong enough to inhibit expression in our system. Presumably these weaker structures do not cause a sufficient roadblock to the initiation of translation by the ribosome. To confirm that the measured decrease in gene expression was indeed due to reduced translation rather than reduced transcription, we used quantitative PCR (qPCR) to verify that our transcript levels per cell remain the same, despite different 5’UTR hairpin sequences being introduced (Supplementary figure 3).

Using the gene expression data, we fitted a model to predict the gene expression output from the calculated MFE values. As the relationship in (Figure 2A) represents a sigmoidal curve, we modified the equation for logistic growth to produce a new equation that predicts gene expression (as normalised fluorescence) from the MFE of folding of the 5’UTR sequence and the maximum output of the promoter used in the gene expression construct (Figure 2B). The fitted curve does not fall within the 95% confidence interval of all the measured data points, suggesting that some relevant properties remain that are not captured by this model. In the most extreme cases, the predicted expression value is 3-fold over-or underestimated compared to the observed value. This compares favourably to the RBS-calculator, which showed deviations of up to 10-fold when it was first published[8].

To test whether this equation has predictive power, we designed, built and tested four further libraries as before by introducing hairpin sequences with degenerate bases between the GAL1 promoter and the GFP coding region (Figure 2C). These four additional libraries (HL3-6) were designed to have weaker average structure strengths within their distributions compared to HL1 and HL2, so that when all six libraries are combined they cover a range from a weak average MFE of -8.0 kcal/mol (HL6) through to a strong average MFE of -32.2 kcal/mol (HL1). The focus was on average MFE values between -32 and -20 kcal/mol, as higher values were predicted to fall within the plateau region of maximum expression and would therefore not lead to significant diversity within the library. The tight peak at maximum expression in library HL6 confirms this.

For each library, the minimal folding energies of all possible members were calculated by RNAfold and then converted into predicted gene expression levels using the established equation and the *Pmax* value of the GAL1 promoter. The predicted distributions were then compared to experimentally-obtained distributions of green fluorescence measurements for each constructed library of yeast cells. An overview of these comparisons is shown in (Figure 2D) and demonstrates a good qualitative agreement between the model predictions and the resulting experimental data.

Generally, the histograms for the predicted normalised expression match those for the measured normalised GFP expression in both their spread (the range of expression levels) and the position of the peak (the average expression). Some peak-broadening is seen in the flow cytometry data compared to the predictions, likely as small and large cells within the measured population cause intrinsic deviations in the data despite cells having the same relative expression. However for the goal of forward engineering, these libraries and the model-based predictions serve their purpose, allowing a user to design a sequence that predictably alters gene expression from a promoter of known strength.

Interestingly, a notable exception in our ability to predict expression from a designed 5’UTR hairpin sequence occurs if the design encodes a *tetraloop* in the 4 base loop sequence of the hairpin. RNA tetraloop motifs, such as those encoded by the sequence UUCG, are known to significantly stabilise hairpin sequences. In hairpin library designs with tetraloop-encoding bases at the loop region, the measured GFP expression from cells was dramatically reduced compared to the predicted expression from the MFE calculations (Supplementary figure 3). Introducing a tetraloop sequence into an otherwise identical library dramatically reduced expression levels and introduced a severe mismatch between predicted and measured expression profiles. This effect was especially pronounced when the stem sequence directly adjacent to the tetraloop was kept devoid of degenerate bases and thereby perfectly complementary. Until RNA folding models can accurately predict the MFE contribution of tetraloops or until the effect of tetraloop inclusion can be accurately modelled, we recommend avoiding tetraloop-encoding bases within hairpin designs.

### 5’UTR hairpins as modular parts

Having predictably altered the expression of GFP from the GAL1 promoter in *S. cerevisiae* with 5’UTR hairpin libraries, we next sought to show that the approach is modular, i.e. that the introduction of designed structures can predictably alter gene expression when combined with other modular DNA parts, such as different promoters. To demonstrate this we first examined the effect of exchanging the open reading frame (ORF) sequence encoding the protein produced in our constructs. We replaced the GFP-encoding sequence with that encoding the red fluorescent protein (RFP) mRuby2 in the constructs for libraries HL4, HL2 and HL1. The sequence identity between the two ORFs encoding these fluorescent proteins is as little as 11.5%, as identified by a BLAST alignment optimised for somewhat similar sequences (blastn) [38].

After cloning the libraries into *S. cerevisiae* we measured the red flourescence per cell of the pooled yeast colonies after galactose induction using flow cytometry and compared the fluorescence distributions to those predicted by our mathematical model (Figure 3A). As seen previously with GFP, the RFP data closely matched the predicted expression levels for all three libraries tested, demonstrating that the 5’UTR libraries remain predictable when the context of the downstream sequence is changed.

**Figure 3:**
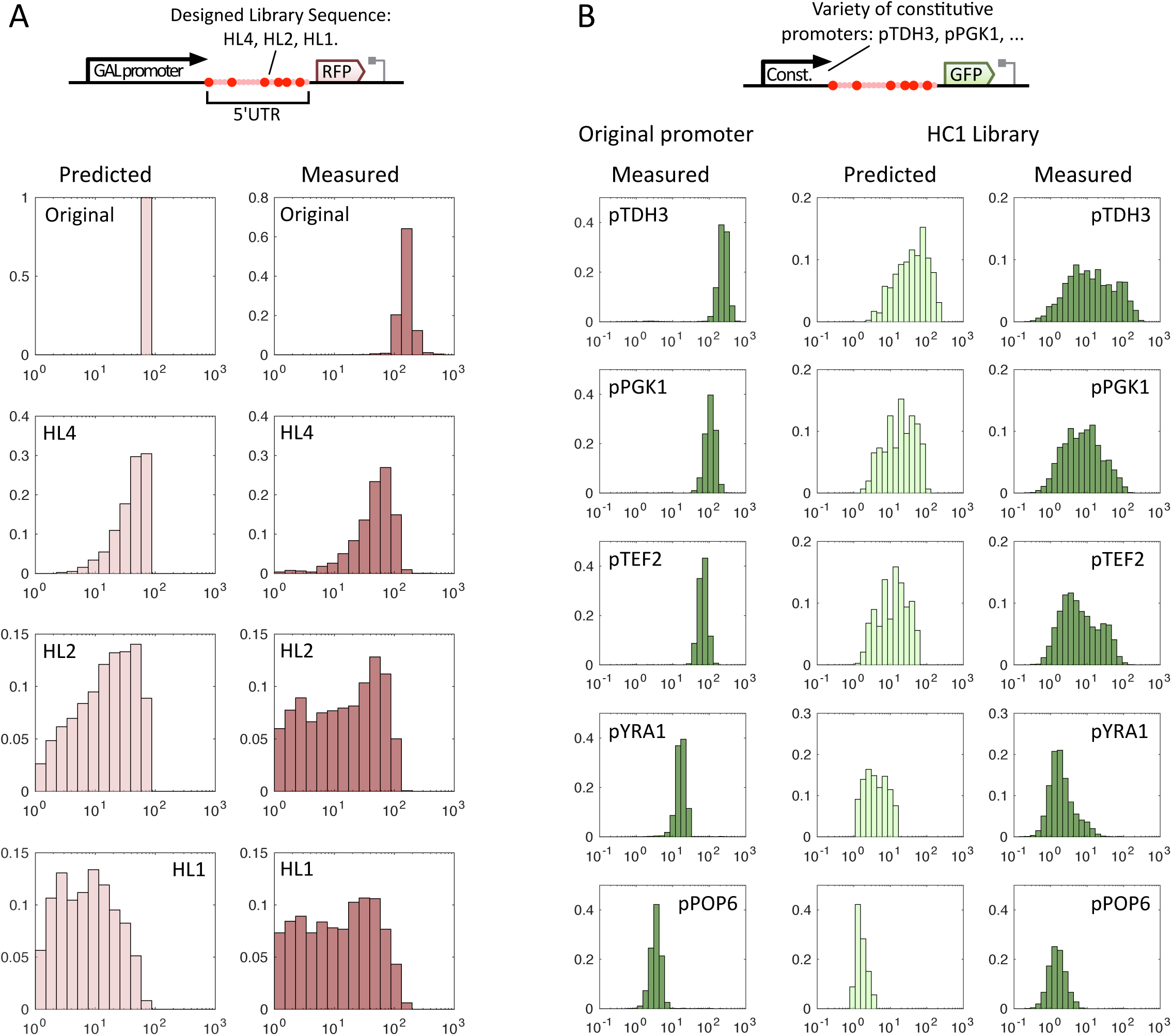
Robustness of predictions with respect to upstream and downstream sequence. Distributions are shown as histograms with normalised fluorescence logarithmically on the x-axis and relative frequency linearly on the y-axis, both unit-less quantities. MFE stands for minimum free energy of folding, a measure of the strength of the hairpin structure in the 5’UTR. (**A**) Robustness of the library predictions to changes in the downstream ORF. The HL4, HL2 and HL1 libraries previously tested with yEGFP are shown here with the mRuby2 ORF. The unmodified GAL1-based promoter is shown as a reference in the top rown. MFE indicates the average MFE of the corresponding library. (**B**) Robustness of the library predictions to the use of different promoters upstream of the 5’UTR hairpin. The HC1 library consisting of hairpins with an average MFE of -28.9 kcal/mol in the 5’UTR for 5 different constitutive promoters of decreasing strength. For comparison, the promoters with a control 5’UTR with no structure are shown in the first column.

Next, we tested the effect of varying the promoters within our constructs. To do this we designed a 5’UTR encoding a new hairpin library with an average MFE of -28.9 kcal/mol. This library, called HC1, was designed to produce a uniform distribution between autofluorescence levels and the maximum expression levels of the various mid-to high-strength promoters. This was cloned upstream of the GFP-encoding ORF and downstream of five constitutive promoters known to have different expression strengths. We then measured the GFP expression from these five promoters in the absence of any 5’UTR structures to obtain *Pmax* values and used these data with the calculated MFE values for the HC1 design to predict the expression profiles of the five new libraries. Histograms of these predictions are shown alongside the equivalent measured data for all five promoters in Figure 3B and show a clear qualitative match.

While this is only a small sample of possible promoters, this result implies that the 5’UTR library approach is modular with respect to upstream promoters (i.e. there is no context-dependency), meaning that the approach could likely be applied to alter the expression from most, if not all promoters in *S. cerevisiae*. However, when the results are inspected with more detail, it is noticeable that the expression level distribution becomes progressively less uniform as weaker promoters are used. For weak promoters, the experimentally measured distribution skews slightly towards lower expression levels.

This phenomenon can be clarified by understanding the use of the logistic fit in Figure 2. An even distribution of expression strengths is only found in the range of values that form the straight part of the curve when the data are plotted against a logarithmic scale (as in Figure 2A). This range becomes narrower for weaker promoters with lower *Pmax* values. Thus, when a 5’UTR library spanning a MFE range of -44 to -24 kcal/mol (i.e. HC1) is used with a weaker promoter, a larger proportion of library members will obtain a sequence that fully-represses detectable expression. This in turn skews the expression level distribution towards the lower end of the spectrum.

However, by simply measuring the normalised fluorescence of a promoter prior to library design and creation, the *Pmax* can be determined and the library MFE spread can be intentionally designed to take into account the promoter strength. Going forward this will allow expression libraries to be created with more precision, regardless of the nature or sequence of the upstream promoter or the downstream ORF.

### Predictable tuning of regulated expression with 5’UTR hairpins

Precise tuning of gene expression is important in many applications of biotechnology, synthetic biology and metabolic engineering. The approach developed here of placing designed secondary structure within the mRNA 5’UTRs offers a new modular tool to achieve this in yeast. Currently in *S. cerevisiae* the most commonly-used method for altering the strength of gene expression is to replace the promoter, and multiple promoter libraries have been described that are intended to enable users to alter the amount of transcription and thus expression of any gene.

Most promoter libraries use sets of constitutive promoters of different strengths, however when gene regulation is required (e.g. for inducible expression) these are not suitable. A small number of promoter libraries based around regulated promoters have been described and used in yeast, but within these libraries the efficiency of regulation often varies considerably as the promoter strengths change. This is because mutating bases within the promoter in order to change the transcription strength often inadvertently alters the efficiency with which transcription factors bind their cognate sites on the promoter and regulate promoter expression.

The 5’UTR hairpin approach developed here is a promising solution to this problem, because the fine-tuning of gene expression output is achieved by altering bases away from the promoter sequence where the regulation is encoded. It therefore offers a new way to modify protein levels from gene expression constructs without affecting their regulation characteristics at the promoter level. To demonstrate this, we directly compared our approach to an equivalent artificially regulated GAL1 promoter library produced previously using targeted mutagenesis (Figure 4).

**Figure 4:**
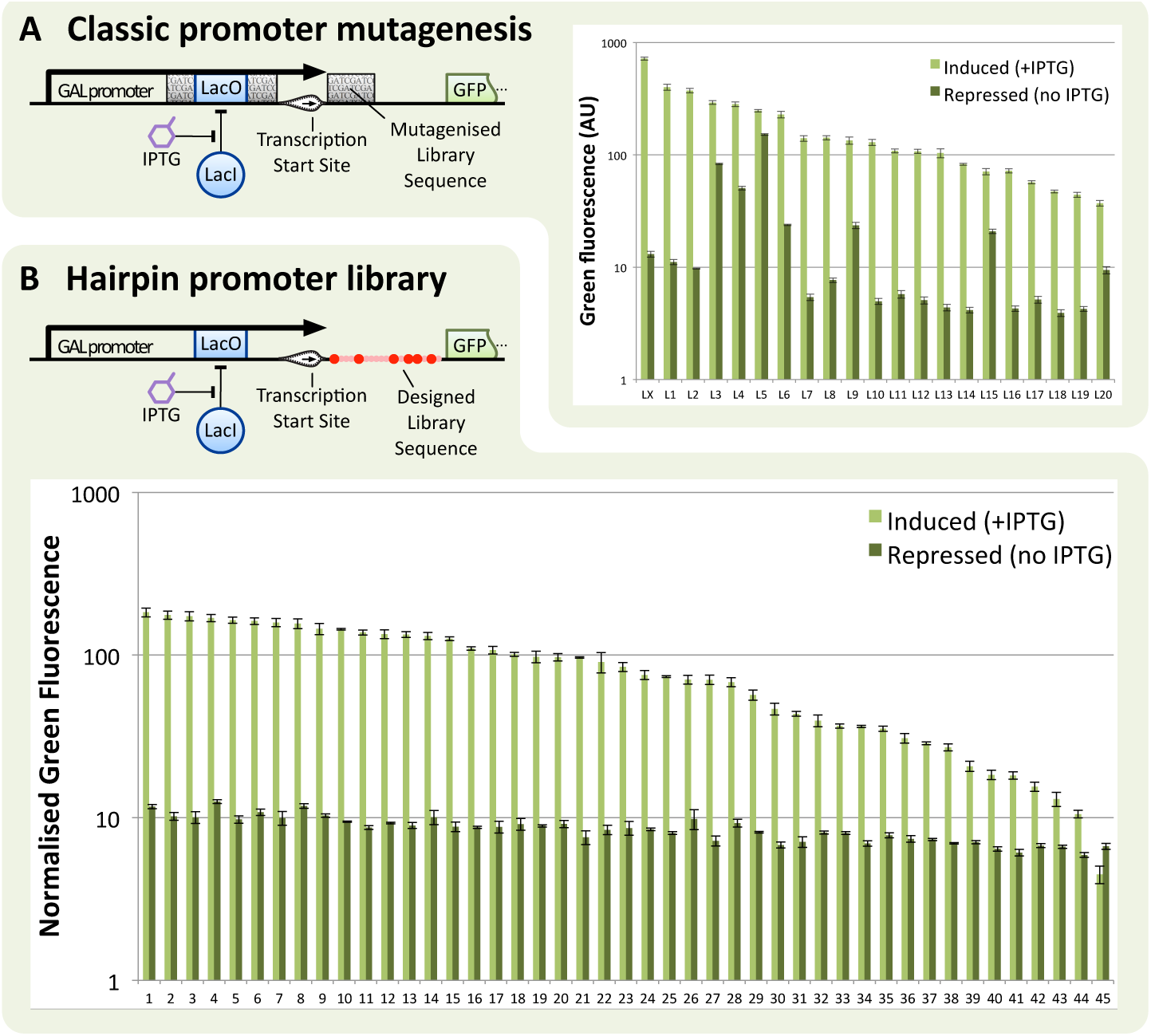
Comparison of methods for regulated expression library creation. The GAL1-based regulated promoters contain a synthetic Lac operator site which can be bound by the Lac Repressor (LacI) in order to repress the promoter. IPTG can subsequently be added to release LacI from the DNA, reversing its repressive effect and inducing yEGFP expression. (**A**) results and method employing targeted random mutagenesis. In this approach, selected regions in the core promoter (grey blocks) are completely randomised. From a pool of 350 candidates, the 20 best performing hits (L1-L20) and the non-mutated version (LX) are selected. Error bars represent standard error of the mean of three biological repeats. (**B**) results and method employing 5’UTR hairpins as developed in this work. A hairpin library sequence (HG1) containing degenerate nucleotides is placed directly following the transcription start site and preceding the start codon. From the resulting clones, 45 are directly picked and characterised, without a prior screening step. Error bars represent standard deviation of the median of three biological repeats.

In both cases, expression of the Lac Inhibitor protein (LacI) in yeast inhibits expression of GFP by binding to an integrated Lac operator site (LacO) within the core of the GAL1 promoter. In galactose media, expression is repressed unless the inducer IPTG is added which blocks the action of LacI and permits full-strength expression.

The 21-member promoter library was previously made by targeted promoter mutagenesis, followed by selection and characterisation experiments requiring a two-week worflow [39]. When measured for GFP expression in induced and repressed conditions (Figure 4A) a desirable range of maximum outputs is seen within the library but the efficiency of repression varies considerably, with several promoters being especially leaky (i.e. not well-repressed).

In contrast, in under a week using our 5’UTR hairpin method we were able to generate a graded library of 45 constructs covering the full range of maximum outputs, while all maintaining strong repression when uninduced (Figure 4B). To do this we simply paired the strongest member of the LacI-repressed GAL1 promoter library (LX) with a library of 5’UTR hairpins (named HG1) designed to have an average folding strength of -28.8 kcal/mol. Upon cloning into *S. cerevisiae* a total of 48 transformants were picked and characterised by flow cytometry for GFP expression in induced and repressed conditions. Only 3 selected colonies with aberrant expression were discarded to yield the 45-member library, whereas the classic promoter mutagenesis method used previously had required screening over 300 colonies to isolate the 21-member graded library.

The use of designed 5’UTR hairpins thus outperforms promoter mutagenesis methods for expression library creation in terms of both ease and speed to create the library and in terms of the resulting constructs showing the desired expression characteristics and no unwanted leaky expression. The increased predictability afforded by this new approach can also aid *a priori* modelling efforts, because there is greatly decreased risk that the promoter with the required maximum expression also has leaky expression or unanticipated impaired regulation. As leakiness typically needs to be as low as possible whenever regulated expression is desired (for example in genetic circuits or biosensors) we anticipate that our 5’UTR hairpin method for tuning expression will be widely-applied.

## Discussion

In this work we have determined how protein expression in *S. cerevisiae* can be predictably tuned through the incorporation of 5’UTR secondary structures. By placing a library of hairpins of different strengths into the 5’UTR region of a gene, we can downregulate translation efficiency and the magnitude of the inhibition can be predicted using well-understood principles of RNA base-pairing. The result is a system for eukaryotes that is similar to the extensively-used RBS Calculator that is available for use in bacteria. We verified the modularity of this system by testing a variety of libraries with two different ORFs and testing one hairpin library with a variety of constitutive promoters. The predictions were shown to hold in all of these cases. In synthetic biology and other disciplines that rely on the precise engineering of gene expression in living cells, the regulation of protein production is of central importance and so being able to fine-tune the efficiency of mRNA translation is an important contribution. Importantly, employing the approach described here requires only a few cloning steps and can be incorporated as a routine part of gene construction and optimisation.

Our results showed that the folding strength of the introduced hairpin determines the expression level of the associated protein with stronger hairpin structures leading to lower expression levels likely through increased interference with the translational machinery of the host during the scanning step of translation initiation. Corroborating this finding, our qPCR experiments showed no difference in the transcript levels when mRNAs contained highly-structured or weakly-structured 5’UTRs, and yet these mRNAs expressed protein at greatly different levels.

Our calibration curve for the mathematical model determines how the folding strength of the hairpin correspond with the translation of the protein and is to our knowledge the most detailed report of this relationship to date. It shows that a MFE difference of 10 kcal/mol is required to impart a 10-fold difference in gene expression, which is more than 7 times the energy that would be expected simply from thermodynamic predictions of the energy required to change the folded to unfolded ratio by 10-fold [40]. This finding is strong evidence that RNA helicase activity in translation initiation partially counteracts the effects of hairpin structure in mRNA. The shape of the curve is effectively a function of the properties of the translation machinery of the host, in particular the processivity of the RNA helicases, notably eIF4A, which are associated with the 40S ribosomal subunit during translation initiation [16, 41].

Furthermore, we also observed that expression was not affected by hairpin structures weaker than -18 kcal/mol. This indicates the translation initiation machinery and associated RNA helicases are likely to be able to fully-denature structures of these strengths. However, hairpins stronger than -44 kcal/mol were not translated at all, suggesting that the native machinery is completely incapable of unfolding these. Precisely determining the limits of the native machinery is important, as it opens up the possibility for determining the impact of individual components of the translation machinery affecting helicase activity, such as eIF4A, eIF4B, eIF4G. Each of these have been shown to play a role in RNA-helicase processivity and observing how overexpression and elimination of these components affect the limits and shape of the curve may provide valuable insights to their function[42, 43, 44].

Interestingly, it was found that weak promoters are especially sensitive to strong 5’UTR structure and thus require appropriate design of libraries with weaker average folding strengths. Another unexpected outcome was the severe impact of tetraloops on translation inhibition. Tetraloops combined with a perfectly paired stem exhibited a far greater inhibitive strength than their net MFE contribution would suggest. While this finding requires more investigation, one possible explanation would be if these motifs were bound and stabilised by other local factors present during translation initiation. Tetraloops such as those tested here, are known to be conserved folding motifs found in ribosomal RNA, which could lead to them being bound by ribosome-associated proteins[45, 46].

Tetraloops are one example of motif that challenges the predictive power of our approach, and other, as yet unidentified features may exist that also affect predictability. Features that attract RNA binding proteins, target the mRNA for degradation or interfere with transport through the nuclear pore are also possibilities. Indeed, in our measurements for the calibration curve, we noted that expression can deviate by up to three-fold from the predicted value in certain cases. This may be caused by the accidental inclusion of sequences encoding unknown motifs and would affect accurate predictions if trying to achieve a single 5’UTRs for a set expression level. For this reason, we prefer the library approach with degenerate bases, which will almost always yield at least one yeast colony exhibiting the desired expression level. As a workaround, hairpins from the pre-characterised list of 5’UTRs shown in Supplementary table 6 can be chosen instead. A further limitation of our approach is that our sequences do not increase expression and only lower it from what is seen normally with the promoter of choice. In bacteria, changing the RBS sequence can often be used to increase expression, however, as yeast 5’UTRs rarely contain rate-limiting secondary structures, the introduction hairpin motifs only reduces the expression levels.

Adapting hairpin design to incorporate active regulation is a promising route for future work. Previous work has shown that aptamer motifs can be designed into a 5’UTR to fold into inhibitory hairpin structures when they bind a specific inducer molecule translation [47]. Approaches such as riboswitches or recruiting RNA-binding proteins such as MS2 coat protein are other routes to regulation. Another interesting potential improvement would be to increase the prediction accuracy by taking into account the dynamic properties of RNA folding. mRNAs with 5’UTR structures have been shown to have biphasic polysome distributions in yeast[19], which indicates that once the structure is unwound it stays unwound, as it reforms slowly relative to the rate of translation initiation. Taking (re-)folding speed into account could therefore lead to improvements in prediction accuracy as has recently been done for the bacterial RBS Calculator [48].

A further improvement to our system would be to make the developed method applicable in all eukaryotes. A new calibration curve will need to be made for each organism that this method is implemented in, since the cellular machinery will behave slightly differently in each case due to the properties of the translation initiation machinery. Interestingly, for higher eukaryotes, such as plants and mammalian cells, the ideal placement of the hairpin module within the 5’UTR may not be the same as in yeast. For work in *S. cerevisiae* we recommend placing the hairpin just upstream of the AUG, as the further upstream it lies, the more likely the encoding-sequence could influence the upstream promoter. However, in higher eukaryotes, the amount of inhibition of translation is thought to be highest when secondary structures are closest to the 5’ cap [18, 49, 19, 21]. This may be because higher eukaryotes have the DHX29 RNA helicase that is absent in *S. cerevisiae* and this is thought to enable them to tolerate much longer, structured 5’UTRs [20].

In terms of applications, we anticipate that this approach will be useful broadly but especially in synthetic biology where exploring and tuning gene expression strength is critical. Already a study on gene expression noise in genetic circuits in *S. cerevisiae* has demonstrated the use of 5’UTR hairpins to modify translation rates of a transcript [50]. We expect our hairpin library system to be ideal for optimising the simultaneous expression of multiple genes. Efforts to create heterologous metabolic pathways in yeast are common in synthetic biology and metabolic engineering [51] and efficient ways of optimising enzyme expression levels in these pathways are needed. Because the fraction of functional library members in a transformed population using our method is high, we expect that multiple libraries can be inserted simultaneously during cloning of a pathway, while at the same time ensuring that functional expression of each gene always occurs.

Due to the different underlying principles of translation initiation in eukaryotes, there are some differences between the approach we present here and the popular RBS Calculator tool for bacteria. In a typical case, the RBS Calculator can be used to both up-and down-regulate, while yeast hairpins only downregulate. However, modularity for bacterial RBS sequences is poor, usually requiring a custom sequence to be designed for each gene. But because hairpin interactions are more specific and predictable this is much less of an issue in our approach, allowing the designed sequences to be used as modules. Already a set of 16 hairpin modules for *S. cerevisiae* 3’UTR regions has been described which modifies gene expression by altering mRNA degradation rates [31]. However, this complementary approach does not allow *a priori* predictions of protein expression levels based on sequence as achieved here.

Taken together, we have developed here a novel sequence-to-output design strategy for the creation of yeast gene expression libraries, and our approach significantly outperforms existing protocols for library creation both in terms of predictability and the speed and ease of the method. With a one-week turnaround time and a maximum of three days of cloning, the use of computationally-designed 5’UTR hairpin libraries is a fast, cheap and accessible way to tune expression levels and to specifically alter the rate of the translation step in eukaryotic systems.

## Acknowledgements

The authors thank Charlie Gilbert for providing critical feedback during the manuscript drafting stage and for Robert Chen for help with the YTK system.

## Supplementary material

### Protocol for the design and construction of 5’UTR hairpin libraries for expression tuning

The cloning process for the creation of 5’UTR hairpin libraries consists of a small number of simple and rapid cloning steps. The cloning strategy was designed to take 3 days or less for DNA construct preparation or up to 7 days from primer delivery to fully characterised library. The simplicity and efficiency of this method are in large part afforded by use of the Yeast Toolkit (YTK) system and by extension Golden Gate cloning [1]. The few accessory plasmids needed in the cloning process can quickly be generated using the collection of parts included in the YTK.

Figure S1 illustrates the process of plasmid construction in detail. The degenerate hairpin sequence is introduced into the 5’UTR by PCR. The primers used in this study, along with the specific libraries they were used for, are shown in Supplementary table S4. The template is a YTK cassette-level plasmid containing the required promoter and ORF in a transcription unit. Template cassettes used in this study are shown in Supplementary table S2. The PCR product is treated with DpnI to remove the template and subsequently circularised in a Golden Gate reaction with BsaI.

To limit the number of cloning steps and the associated loss in library diversity, the self-ligated PCR product is used directly in the multigene assembly step without passage through *E. coli*, as is customary in the YTK protocol. Equimolar quantities of the product are used in a multigene assembly step, with a pre-assembled yeast integration cassette and a cassette for constitutive mRuby2 expression. The presence of red fluorescent protein is later used as a control for correct plasmid assembly. Consequently, correct clones do not have to be hand-picked during the cloning process, which is unworkable in a library approach with many thousands of individual library members. A list of all libraries created in this study and which cassettes were used in their respective multigene assemblies is provided in Supplementary table S1.

To speed up the process of cloning and to maximise library diversity, the multigene assembly reaction is transformed into the fast growing NEB Turbo competent *E. coli* strain. To allow for parallelization, the transformation is performed with chemically competent cells. The turbo strain can be grown up for miniprep in as little as 5 hours, allowing considerable time savings. To complete the library construction process, the pooled and miniprepped multigene assemblies are digested with NotI in preparation for yeast transformation. The transformation itself is carried out using a high efficiency protocol [2] and a large amount of linearised DNA (3-5 *μ*g) to typically yield thousands of library candidates.

Prior to wet-lab activities, the desired libraries are designed in-silico. Library design starts with the selection of a scaffold hairpin, which represents the strongest structure in the hairpin library space (i.e. the strongest folding library member). The scaffold hairpin must conform to the following design requirements:

- The sequence is entirely or largely palindromic (it is a strong and perfect hairpin).
- The sequence has a minimum free energy of folding that is as low or lower as the lowest required member of the library.
- There is no premature start codon contained in this sequence.
- The loop of the hairpin does not contain an unusually stable sequence known as a tetraloop (see below).
- The two halves of the hairpin are short enough to be contained in the tails of primers used to amplify the selected transcription unit cassette.
- The sequence does not contain BsaI, BsmBI or NotI restriction sites.
- The sequence does not contain other forbidden restriction sites or sequences that interfere with the function of the construct in the chosen application.

In our experiments, the UUCG tetraloop was shown to have a dramatic and unpredictable effect on expression. Inclusion of tetraloop sequences such as GNRA [3, 4], UNCG [3, 5], CUYG [3, 6], UNAC [7] and ANYA [8] is therefore advised against.

Initially the hairpin scaffolds used in this project were based on the ideal Lac operator sequence, which is palindromic. These were subsequently extended and modified to accommodate stronger libraries and shorter cloning primers. However newly designed scaffold can be based on the native 5’ UTR -by adding an inverted repeat of this sequence -or on a de-novo structure created through one of many inverse RNA structure prediction solutions available [9, 10, 11].

When a scaffold hairpin has been selected, the degeneracies can be inserted at any base paired position in the hairpin. Substitution of the cytosine in a G-C base pair is preferred, as the opposing guanine is capable of forming the weaker non-canonical G-U pair in addition to the strong canonical pair. This allows variety to be introduced into the hairpin without disrupting its basic structure. In terms of degenerate nucleotide code this is the substitution of the cytosine in a G-C base pair by degenerate nucleotide Y (i.e. C or T). More variety can be created still by also allowing a non-pairing base at this position with degenerate base H (C, T or A). Inclusion of guanine at a locations where it was not originally present is generally avoided because of its large potential for (unintended) interactions. For the same reason, W is the only degenerate nucleotide generally substituted at an A-T pair. Additionally, degeneracies introduced at both sides of the pair are generally avoided, as the probability for a matching basepair decreases drastically as the number of possibilities at each side of the pair increases.

With the scaffold hairpin and degeneracies set, a location in the 5’UTR must be chosen for its insertion. To ensure modularity and conservation of the Kozak sequence, the hairpin is inserted 5-15 bp upstream of the ATG. Further upstream may reduce its effect [12]. To keep the total length of the 5’UTR similar to native 5’UTRs and to ensure the hairpin is not disrupted by unintended basepairing with upstream sequence, the 5’ end of the hairpin must be within 10 bp of the transcription start site.

For the next step in the process a fasta file is generated containing all possible sequences that can arise from the selected degeneracies. There are indications that secondary structure involving the start codon has a strong impact on repression. We therefore chose to include the sequence up to that point in the calculations. The cutoff directly after the ATG is still somewhat arbitrary and may be optimised in the future.

The generated list of sequences in multi-FASTA format serves as the input for the RNAfold script used for the folding energy estimation. This is part of the ViennaRNA package 2.0, which is a widely used suite of tools centred around the many facets of RNA structure analysis and prediction [11]. The key non-default parameter used in this script is -T30. This modifies the energy parameters used in the script to simulate 30°C, the typical temperature for yeast growth. The full command for script execution is:

~~~
RNAfold -T30 -d2 --noLP --noPS *<* Sequence list 1.fa *>* Output MFE list 1.txt
~~~

The MFE value for each of the input sequences is then extracted from the output file. This list is used to create a histogram for visualisation of the distribution of folding energies and further to calculate the predicted distribution of expression levels. The logistic fit obtained in Figure 2 in the manuscript is used to convert the MFE values to the predicted normalised median fluorescence value for each of the members of the library. Finally, the obtained expression values are plotted as a histogram and compared to the intended expression profile. At this point adjustments can be made to the degeneracies in the hairpin scaffold until the predicted outcome matches the desired outcome.

This pipeline has been implemented in a python script. As input, this script takes the degenerate library sequence and it outputs the associated histograms for the predicted MFE and normalised expression values. It can also output a list of all library member sequences and their associated MFE and expression values. The command for script execution to generate histograms and a complete list of all members for the HL1 library is:

~~~
python degFoldPlotMod.py -T 30 --prom strength 210 --output tsv HL1 AGTATCAACAAAAAgaattgtgagYSYtYatca gagcgctYaYaattYttTTYgtYGagYAGAAAAACCCCAAATATG
~~~

The script requires the normalised promoter strength of the promoter that is being used - in this case 210 for the GAL1 promoter - and handles the execution of the RNAfold software without requiring further user interaction. The generated sequences are also analysed for inadvertently introduced premature start codons and restriction sites.

**Figure S1:**
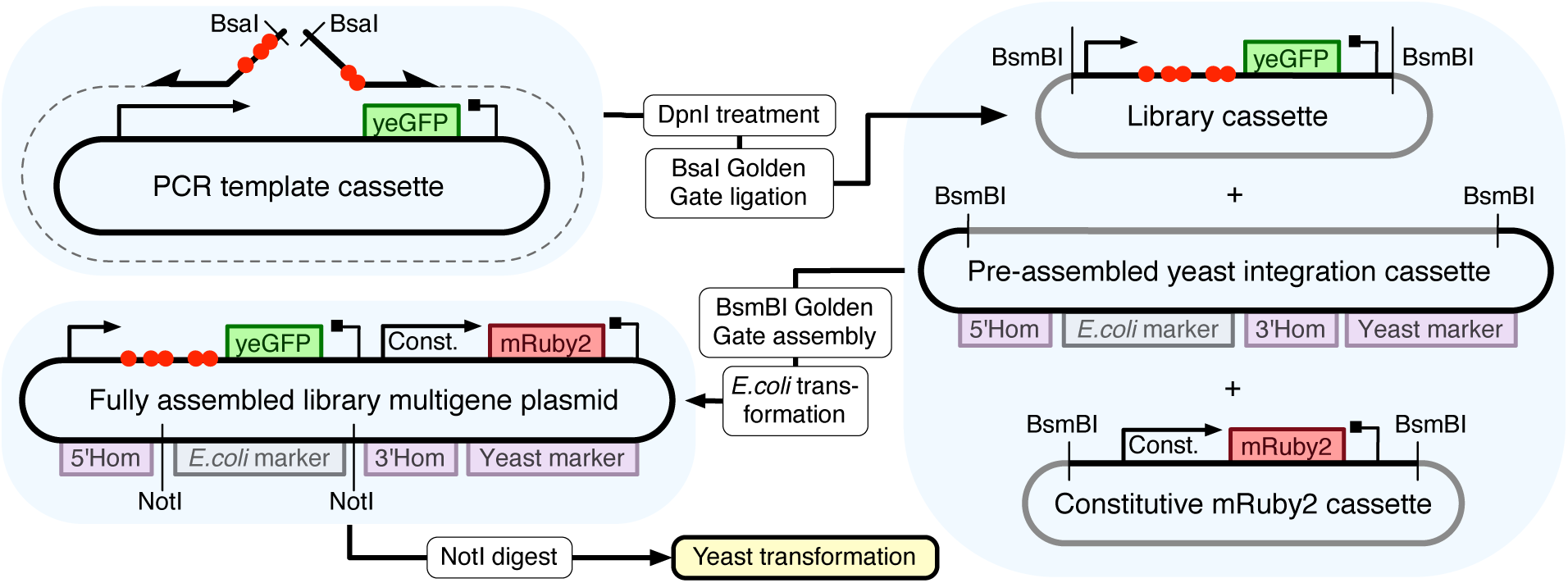
5’UTR hairpin library construction using the Yeast ToolKit (YTK) system. **1)** Primers encoding the hairpin library of choice (indicated by the red spheres) are used to amplify a template cassette carrying the selected promoter and a green fluorescent reporter gene (yEGFP). A DpnI digestion is performed to eliminate the template and the PCR product is subsequently self-ligated in a BsaI Golden Gate reaction. **2)** The resulting plasmid is purified and directly used in the multigene assembly, bypassing the typical intermediate transformation in order to preserve library diversity. The multigene BsmBI Golden Gate assembly is performed with two additional cassettes: a preassembled yeast integra-tion cassette containing a selectable marker and homology regions for integration, and a constitutively-expressing red fluorescent reporter gene cassette (mRuby2). **3)** NEB turbo competent *E. coli* is transformed with the multigene assembly. The thousands of resulting transformants are pooled and miniprepped. Finally the library of plasmids is digested with NotI before being used in a high efficiency yeast transformation resulting in hundreds to thousands of 5’UTR library candidates.

**Figure S2:**
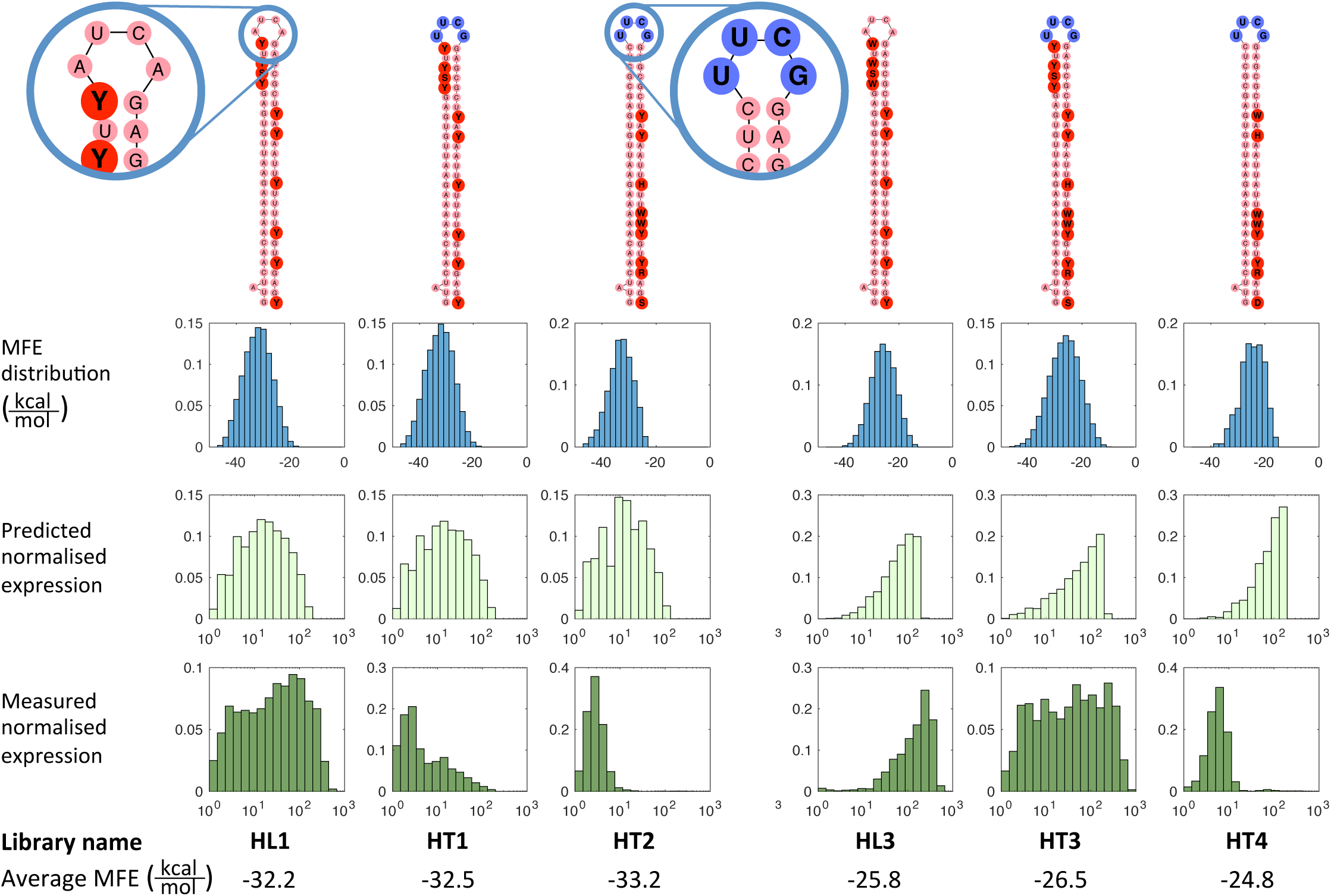
Two sets of libraries characterising the effect of the highly stable tetraloop on fluorescence levels. Left set (first three columns) each have MFE averages of -32 to -33 kcal/mol. The right set has MFE averages of -24.5 to -26.5 kcal/mol, as indicated below the histograms. The illustrations at the top show the scaffold hairpin and introduced degeneracies for each library. The blue bases in the RNA structures indicate the presence of the strong UUCG tetraloop. The top row of normalised histograms shows the MFE distributions for the 5’UTRs of each of the 6 libraries. The horizontal axes for these panels ranges on a linear scale from -50 kcal/mol on the left to 0 kcal/mol on the right. The middle row converts these into a distribution of predicted expression levels. Expression profiles are predicted by mapping the structure MFE to an associated expression level using the function derived in **Figure 2A and B**. The third row shows the distribution of fluorescence levels in yeast cells as measured by flow cytometry. In the lower two rows, the horizontal axis corresponds to normalised green fluorescence (a unit-less quantity) ranging on a logarithmic scale from 1 on the left to 1000 on the right. Note that measured expression levels deviate substantially from the predictions for libraries that contain a tetraloop. Note also that libraries HL1 and HT1 are identical apart from the loop sequence, that libraries HT2 and HT4 have no degeneracies in the section of the stem that is directly adjacent to the tetraloop sequence and the dramatic effect these properties have on the measured expression.

**Figure S3:**
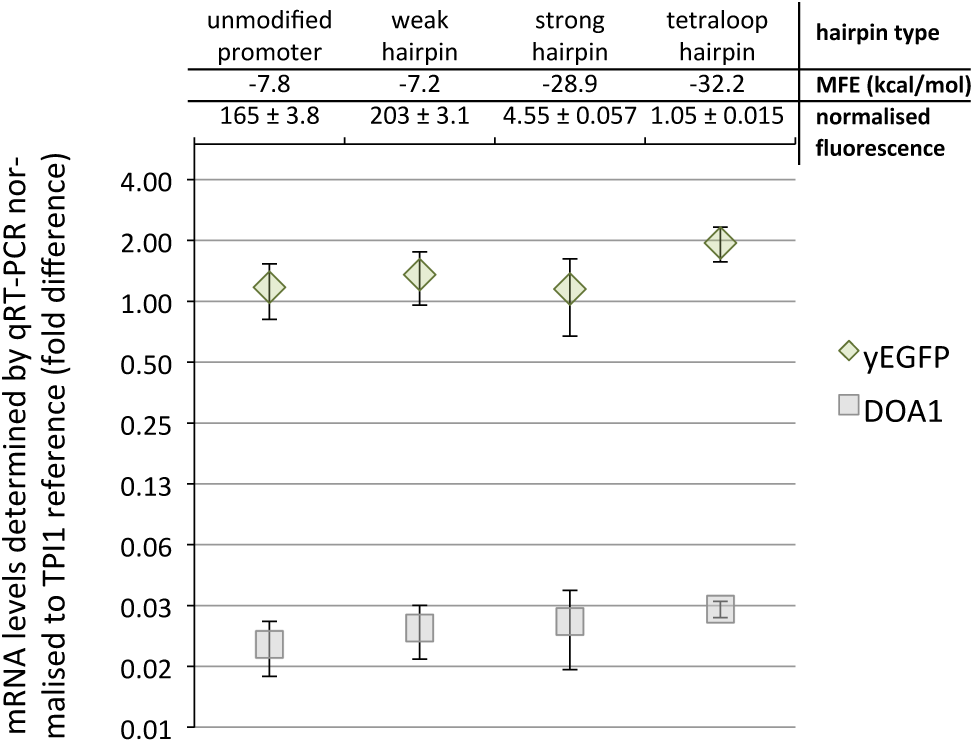
Low impact of 5’UTR structure on transcript levels per cell. Two-step qPCR was used to determine transcript levels in strains with hairpins of various strengths in the 5’UTR of yEGFP expressed from a pLX GAL1-derived promoter. Associated folding energies (Minimum Free Energy, MFE in kcal/mol) and normalised expression strength of each construct is shown at the top of the graph. Note that mRNA levels are not affected by changes in expression strength between the constructs. Expression strengths are normalised against cellular autofluorescence of the parental strain. Transcript levels are normalised against the TPI1 mRNA, using the dd-Ct method [13]. TPI1 is a strongly expressed reference gene (˜ 200 mRNAs per cell), while DOA1 is a weakly expressed reference gene (2.6 mRNAs per cell on average). Error bars indicate standard deviation of technical triplicates.

**Table S1:**
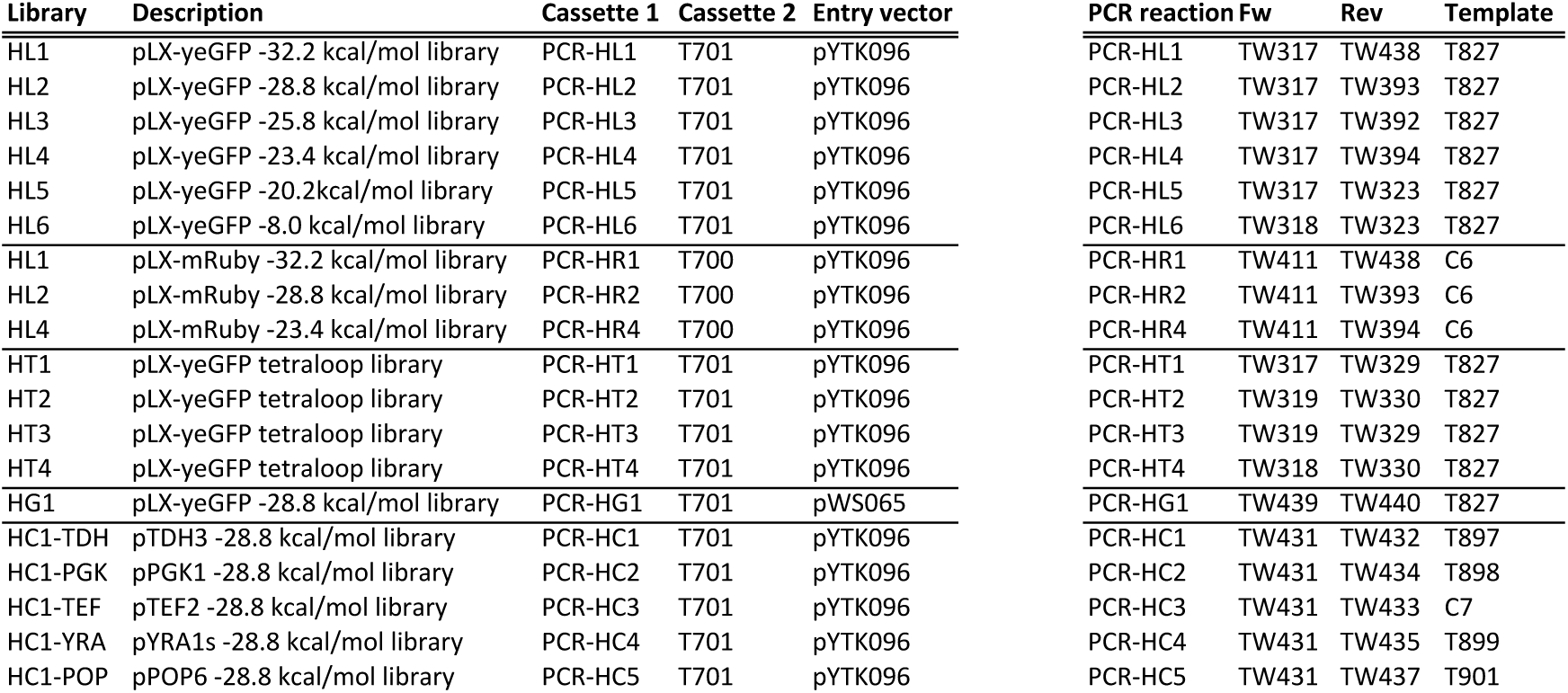
Yeast ToolKit multigene level library plasmids with corresponding PCR-generated cassette-level parts. Contrary to normal YTK protocol, one of the cassette-level parts is generated directly with PCR. PCR amplifications were subjected to DpnI treatment and a BsaI-Golden Gate digestion-ligation to circularise the product prior to incorporation into the multigene YTK assembly. Cloned cassette plasmids and template plasmids used in the PCR reactions are detailed in Supplementary table S2.

**Table S2:** Yeast ToolKit cassette-level plasmids used in the construction process, as template for PCR or directly in a multigene-assembly step. Part-level plasmids used in the assembly of these cassettes are listed. Sequences for part-level plasmids not included in the YTK are given in Supplementary table S3.

| Name | Description | Parts used in assembly |  |  |  |  |  |  |
| --- | --- | --- | --- | --- | --- | --- | --- | --- |
| T827 | pGAL1-yEGFP cassette | pYTK002 | T655 | T675 | pTMP065 | pYTK068 | pYTK095 |  |
| C6 | pGAL1-mRuby2 cassette | pYTK002 | T655 | pYTK034 | pTMP065 | pYTK068 | pYTK095 |  |
| T897 | pTDH3-yEGFP cassette | pYTK002 | pYTK009 | T675 | pTMP065 | pYTK068 | pYTK095 |  |
| C7 | pTEF2-yEGFP cassette | pYTK002 | pYTK014 | T675 | pTMP065 | pYTK068 | pYTK095 |  |
| T898 | pPGK1-yEGFP cassette | pYTK002 | pTMP030 | T675 | pTMP065 | pYTK068 | pYTK095 |  |
| T899 | pYRA1s-yEGFP cassette | pYTK002 | pYTK018 | T675 | pTMP065 | pYTK068 | pYTK095 |  |
| T901 | pPOP6-yEGFP cassette | pYTK002 | pYTK024 | T675 | pTMP065 | pYTK068 | pYTK095 |  |
| T700 | pTEF-yEGFP cassette | pYTK004 | pYTK013 | T675 | pYTK054 | pYTK072 | pYTK095 |  |
| T701 | pTEF-mRuby2 cassette | pYTK004 | pYTK013 | pYTK034 | pYTK054 | pYTK072 | pYTK095 |  |
| pWS065 | pre-assembled HIS integration cassette | pYTK008 | pYTK047 | pYTK094 | pYTK073 | pYTK076 | pYTK088 | pYTK090 |

**Table S3:**
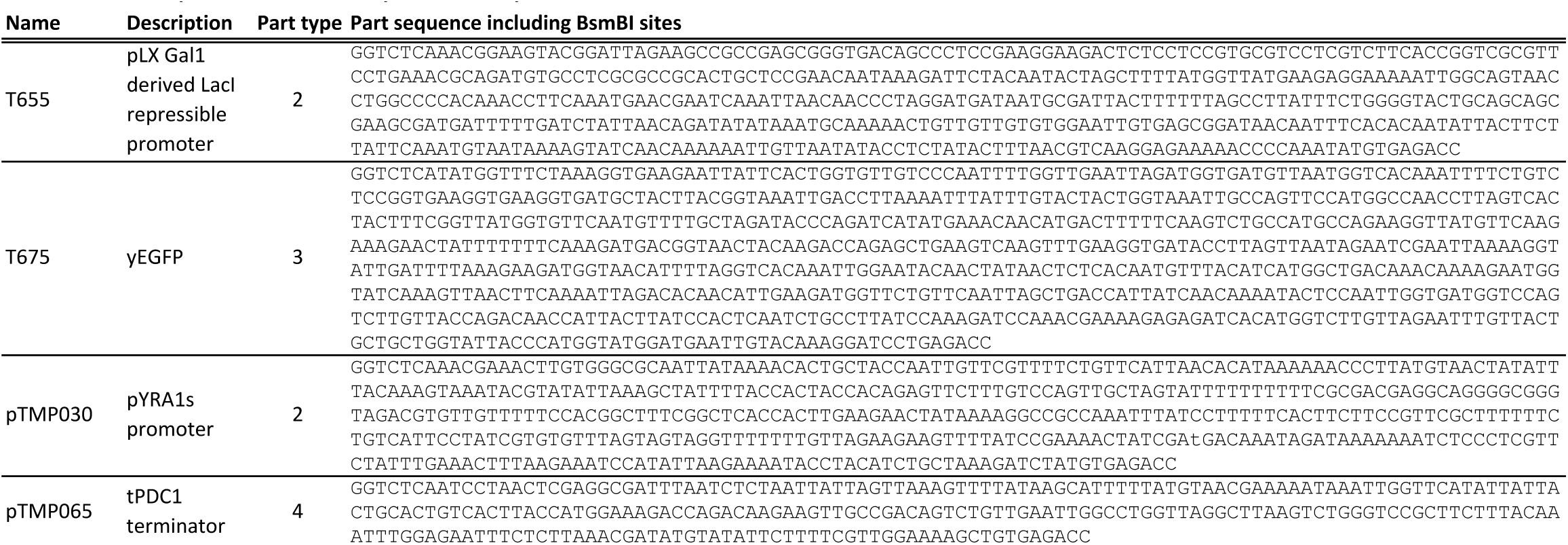
Custom part-level plasmids for the Yeast ToolKit used for the creation of 5’UTR hairpin libraries. The standard backbone sequence for YTK part-level plasmids is not shown.

**Table S4:**
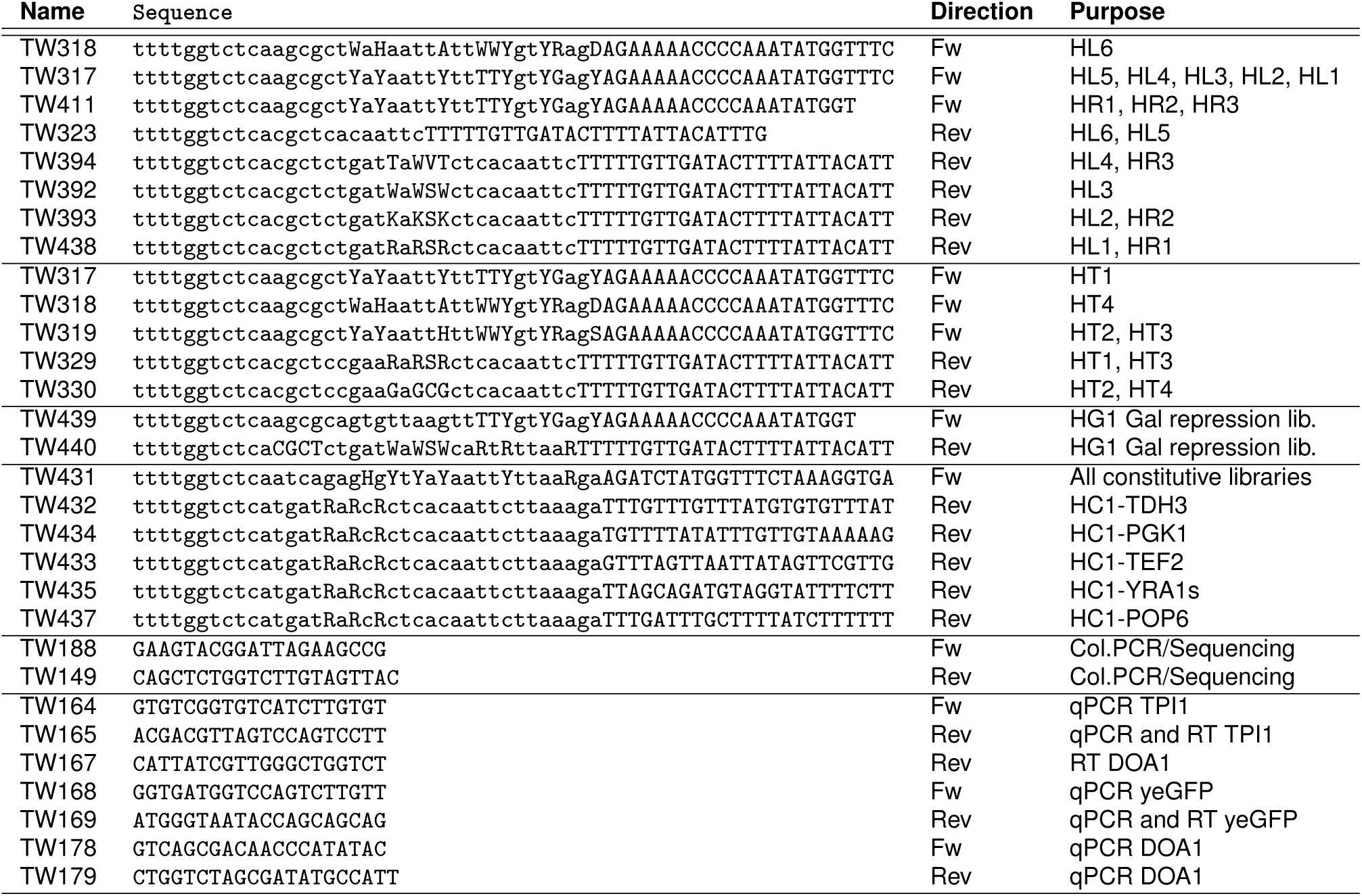
Primers used in this study. Capital letters indicate annealing sequence and locations of nucleotide degeneracies. W = A or T; H = A, C or T; Y = C or T; R = A or G; D = A, G or T; V = A, C or G; S = G or C; K = G or T

**Table S5:**
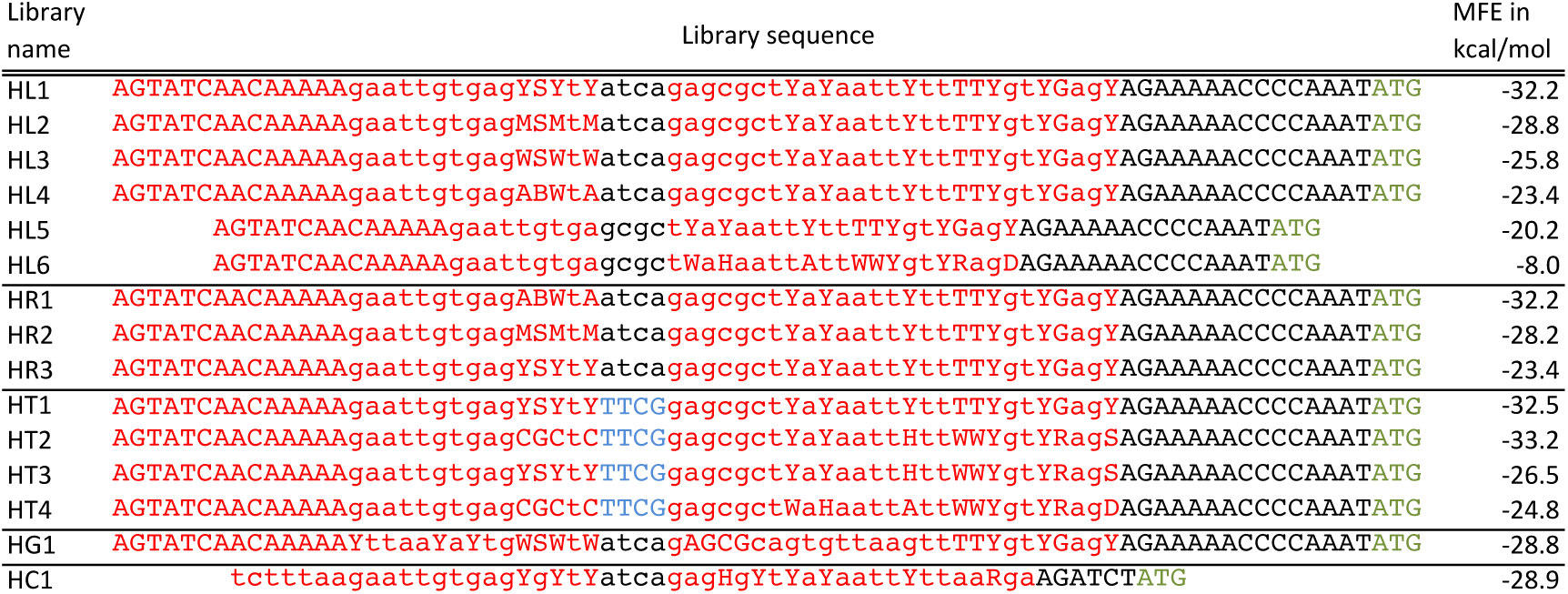
Sequences of the various hairpin libraries tested in this study. Hairpin stems are annotated in red, tetraloops in blue and the start codon in green. MFE in kcal/mol indicates the average minimum free energy of folding of each of the hairpin libraries.

**Table S6:**
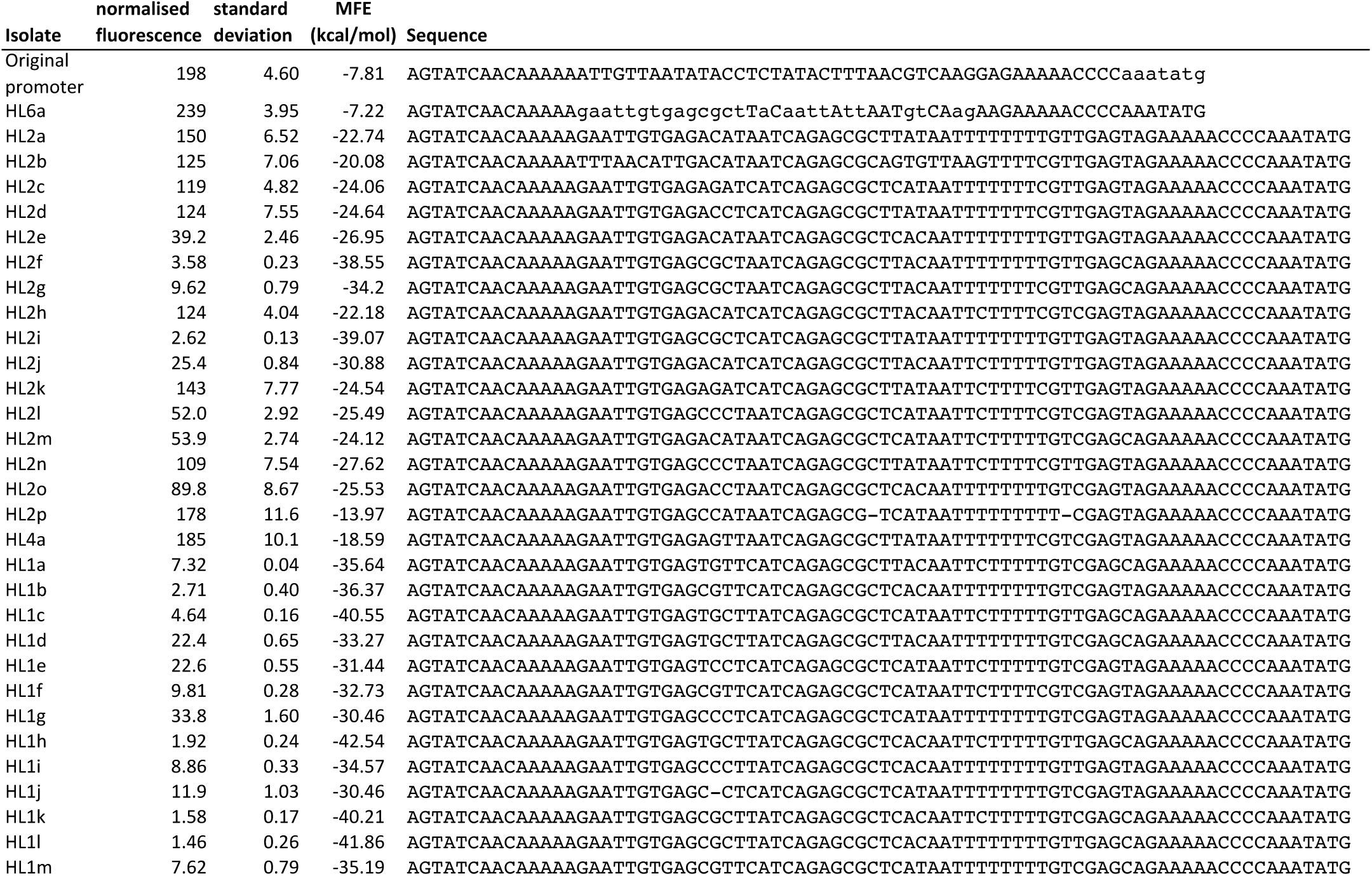
List of all library isolates displayed in figure 2A. For each isolate, the following properties are listed: parental library, normalised median fluorescence, standard deviation of the fluorescence over three biological replicates, predicted minimum free energy of folding of the 5’UTR and its DNA sequence.

